# Peptide:MHC dependent activation of natural killer cells through KIR2DS2 generates anti-tumor responses

**DOI:** 10.1101/2020.04.15.042077

**Authors:** Pauline Rettman, Matthew D. Blunt, Berenice Mbiribindi, Rebecca Fulton, Ralf B. Schittenhelm, Andres Vallejo Pulido, Leidy Bastidas-Legarda, Marta E. Polak, Rochelle Ayala, Anthony W. Purcell, Aymen Al-Shamkhani, Christelle Retiere, Salim I. Khakoo

**Affiliations:** School of Clinical and Experimental Sciences, Faculty of Medicine, University of Southampton, Southampton, UK; Monash Biomedical Proteomics & Metabolomics Facility, Biomedicine Discovery Institute and Department of Biochemistry and Molecular Biology, Monash University, Clayton, Victoria, Australia; Biomedicine Discovery Institute and Department of Biochemistry and Molecular Biology, Monash University, Clayton, Victoria, Australia; School of Cancer Sciences, Faculty of Medicine, University of Southampton, Southampton, UK; Etablissement Français du Sang, 44011 Nantes Cedex 01, France

**Author notes:** These authors contributed equally to this work (Pauline Rettman, Matthew D. Blunt). Department of Surgery/Transplant Division, Stanford University School of Medicine, 1201 Welch Road, MSLS P328, Stanford, CA 94305-5492. To whom correspondence should be addressed: Salim I. Khakoo, Faculty of Medicine, University of Southampton, Henry Wellcome Laboratories, Level E, South Academic Block, Southampton General Hospital, Tremona Road, Southampton, SO16 6YD. Author contributions: SIK, PR, MDB, BM, RBS designed experiments; PR, MDB, BM, RF RBS, LBL, RA performed experiments; AVP, PR, MDB, RBS and SIK analyzed data; CR provided key reagents; SIK, AA-S, AWP, MP supervised experiments and interpreted data; PR, MDB, RBS, AVP, AA-S and SIK wrote the manuscript.

**Keywords:** Natural Killer cells, Killer cell immunoglobulin-like receptors, MHC, vaccination

## Abstract

Natural killer (NK) cells are key components of the immune response to viral infections and cancer. Their functions are controlled by activating and inhibitory killer-cell immunoglobulin-like receptors (KIR) which have MHC class I ligands. KIR2DS2 is an activating KIR, that binds conserved viral peptides in the context of HLA-C and has been associated with protective responses to both cancer and viral infections. We sought to investigate whether NK cells can be specifically activated in a peptide:MHC dependent manner to generate functional immune responses as a potential immunotherapeutic strategy.

We developed a peptide-based KIR targeting DNA vaccine. Immunizing KIR-Tg mice with the vaccine construct generated *in vivo* peptide-specific activation of KIR2DS2-positive NK cells leading to canonical and cross-reactive peptide specific immune responses *in vitro*, and also *in vivo* inhibition of tumor growth. Using immunopeptidomics we identified that the nuclear export protein XPO1, which has been associated with a poor prognosis in many different human cancers, furnishes an HLA-C restricted cancer-associated peptide ligand for KIR2DS2-positive NK cells. We thus define a novel strategy to activate KIR in a peptide-specific manner and identify a rationale for its use in cancer immunotherapy.

**Significance statement:** Natural killer (NK) cells are known to have important roles in determining the outcomes of viral infections and cancer. The killer cell immunoglobulin-like receptors (KIR), and in particular the activating receptor KIR2DS2, have been associated with the outcome of a number of different human cancers. Specific activation of NK cells through KIR2DS2 is challenging because it shares high (>98%) sequence homology with related inhibitory KIR. We have used a peptide:MHC targeting strategy to activate NK cells through KIR2DS2 and identified a novel cancer-associated ligand for this receptor. The work provides a proof-of-concept for targeting NK cells through activating KIR as a cancer immunotherapy strategy.

## Introduction

The potential of natural killer (NK) cells for treating cancer is now becoming recognized (1). However defining the role of NK cells in a personalized medicine strategy to treat cancer requires detailed understanding of the receptor:ligand interactions between NK cells and their cancer targets (2, 3). One important family of NK cell receptors are the killer cell immunoglobulins like receptors (KIR). These form a polymorphic gene family of receptors with MHC class I ligands (4). KIR have been implicated in susceptibility to and the outcome of many different cancers (5-11). Thus targeting the KIR has the potential to form part of a therapeutic strategy to treat cancer (12).

The KIR can be activating or inhibitory with the MHC class I ligand specificities of the inhibitory KIR being relatively well defined. However the ligand specificities of the activating KIR have been much harder to identify. Recent work has shown that activating KIR can have an MHC class I-restricted peptide specificity (13-16). Whilst T cell receptors have a tight restriction on the peptide:MHC complexes that they bind, the KIR recognize families of peptide:MHC complexes in a motif-based manner allowing recognition of multiple peptides and HLA class I allotypes (17, 18). KIR2DS2 is an activating receptor that recognizes group 1 HLA-C molecules in combination with different viral and synthetic peptides, and we have recently shown that KIR2DS2 recognizes highly conserved flaviviruses and hepatitis C peptides with an alanine-threonine sequence at the C-terminal −1 and −2 positions of the peptide in the context of HLA-C (14, 19).

To date no cancer-associated peptides that bind activating KIR have been identified. This is relevant as activating KIR have been associated with protective responses against cancer. KIR haplotypes containing activating KIR confer protection against relapse of acute myeloid leukemia (AML) following bone marrow transplantation, and this has been mapped to the region of the KIR locus that contains KIR2DS2 (20-22). In cord blood transplantation this benefit is accentuated if the recipient of the transplant has group 1 HLA-C allotypes, the putative ligands for KIR2DS2 (23). KIR2DS2 has also been associated with protection against a number of solid tumors including cervical neoplasia, breast cancer, lung cancer, colorectal cancer and hepatocellular carcinoma (6, 7, 24-26). *In vitro*, recognition of cancer cell lines for KIR2DS2 has been observed, but is not specific and also encompasses inhibitory KIR2DL2/3 (27). Additionally, KIR2DS2-positive NK cells appear to express higher levels of FcγRIII (CD16), have enhanced functionality and confer enhanced protection against glioblastoma in a xenograft model (28). Consistent with this enhanced functionality, in a clinical trial of an anti-GD2 antibody in neuroblastoma, KIR2DS2-positive patients had improved survival compared to KIR2DS2-negative patients (29).

Targeting KIR2DS2 in an immunotherapeutic strategy is challenging as it shares more than 98% sequence homology with the inhibitory receptors KIR2DL2 and KIR2DL3. However KIR2DS2 does have a distinct peptide:MHC specificity suggesting that there is potential for developing peptide-based approaches to activate NK cells in a manner analogous to targeting cytotoxic T cells (30). Furthermore as it has a broad peptide:MHC class I specificity, it has the potential to recognize multiple different peptide:MHC combinations, consistent with observations of protection in both viral diseases and cancer (14, 23). To investigate the therapeutic potential of targeting activating KIR in a peptide-dependent manner, we have used a DNA vaccine strategy to activate NK cells through KIR2DS2 and generate anti-tumor immune responses.

## Results

### Activation of NK cells using peptide:MHC mediated ligation of KIR2DS2

To assess the potential for peptide-based activation of NK cells we developed peptide:MHC (pMHC) DNA constructs that expressed HLA-C*0102 linked to a peptide separated by a T2A self-cleaving sequence. Two previously described KIR2DS2-binding peptides were used for these: LNPSVAATL (C*0102-LNP) and IVDLMCHATF (C*0102-IVDL) (14). The constructs were used as naked DNA vaccines to immunize KIR transgenic (KIR-Tg) mice (31). These mice express a human KIR haplotype B locus that includes KIR2DS2 and are backcrossed onto an MHC class I negative background. KIR-Tg mice were injected intramuscularly with 50µg of DNA, without additional adjuvant, weekly for two weeks. Mice were sacrificed one week after the final injection, and splenocytes and hepatic lymphocytes isolated. Results were compared to DNA constructs encoding HLA-C*0102 alone, or DNA encoding HLA-C*0102 in combination with a control peptide, IVDLMCHAAA (C*0102-AAA), in which the P9 and P10 residues of the peptide IVDLMCHATF had been mutated to alanine to abrogate binding to KIR2DS2 and HLA-C*0102 respectively. Injection with DNA encoding viral peptides induced activation of splenic NK cells as indicated by KLRG1 expression, but was greatest in the C*0102-IVDL group (40%) compared to C*0102-AAA (28%), p<0.001 (Fig. 1A and Fig. S1 for gating strategy). However we did not observe a specific increase in the frequency of KIR2DS2-positive NK cells (data not shown), suggesting additional signals are required to drive NK cell proliferation. In paired analyses KLRG1 expression was upregulated to a significantly greater extent on KIR2DS2-positive (KIR2DS2+) versus KIR2DS2-negative (KIR2DS2-) splenic NK cells (p<0.01 for C*0102-IVDL and p<0.05 for C*0102-LNP), but not by control constructs (Fig. 1B). Activation was more marked in mice injected with C*0102-IVDL compared to C*0102-LNP, consistent with the stronger binding *in vitro* noted previously in tetramer binding experiments (14).

**Fig. 1:**
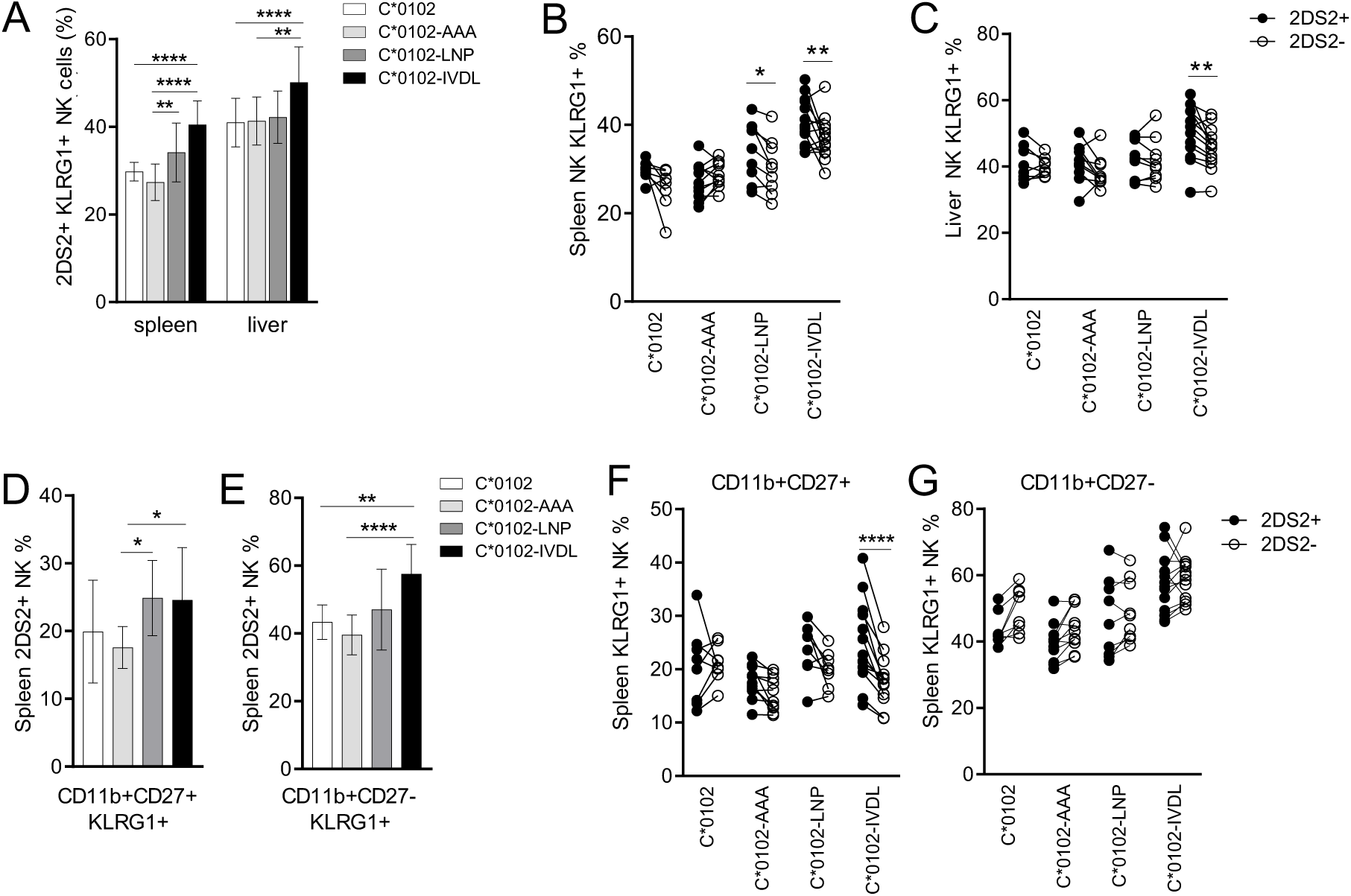
A peptide:MHC DNA vaccine that targets KIR2DS2 activates NK cells. KIR-Tg mice were injected intramuscularly using two doses of the indicated DNA construct one week apart and then assessed for activation of NK cells by flow cytometry for expression of KLRG1. **A)** The frequency of KLRG1 expression on KIR2DS2+ NK cells in the spleen and livers of KIR-Tg mice vaccinated with DNA plasmids containing HLA-C*0102 (C*0102, white bars), HLA-C*0102 plus IVDLMCHATAAA (C*0102-AAA, light gray bars), HLA-C*0102-LNPSVAATL (C*0102-LNP, dark gray bars), HLA-C*0102-IVDLMCHATF (C*0102-IVDL, black bars). **B, C)** Comparison of KLRG1 frequencies on KIR2DS2+ (filled circles) or KIR2DS2- (open circles) CD3-NK1.1+ NK cells in the spleens (**B**) and livers (**C**) following vaccination. **D, E)** KLRG1 on splenic CD11b+CD27+ (**D**) and CD11b+CD27- (**E**) NK cell sub-populations following vaccination. **F**,**G)** Comparison of KLRG1 expression on KIR2DS2+ and KIR2DS2-splenic NK cells in the CD11b+CD27+ (**F**) and CD11b+CD27- (**G**) sub-populations. N=8-14 mice per group. For all plots comparisons between two groups were made by paired t test (2 groups) and 2-way ANOVA (more than 2 groups) (*p < 0.05, **p< 0.01, ****p<0.001).

Furthermore, activation of hepatic NK cells was noted only in experiments using the C*0102-IVDL construct, with the expression of KLRG1 being significantly higher on the KIR2DS2+ versus KIR2DS2-population (p<0.01) (Fig. 1C). KLRG1 was upregulated by C*0102-IVDL vaccination on both mature CD11b+CD27+ and terminally mature CD11b+CD27-splenic NK cells compared to C*0102-AAA: 27% versus 18% (p<0.05) and 62% versus 40% (p<0.001) respectively (Fig. 1D and E). Specific activation of KIR2DS2+ versus KIR2DS2-NK cells was most marked on the CD11b+CD27+ splenocytes (p<0.001), as compared to terminally differentiated CD11b+CD27-NK cells (Fig. 1F and G). Similarly after four weekly injections we observed activation of NK cells, with upregulation of KLRG1 on splenic and hepatic NK cells with both peptide-containing constructs, and KIR2DS2-specific activation in the CD11b+CD27+ double-positive population with C*0102-IVDL (Fig. S2).

NK cells from C*0102-IVDL and C*0102-AAA vaccinated mice were profiled by RNA seq to determine changes in expression one week following vaccination (Fig S3 for gating strategy). Principal component analysis (PCA) showed that KIR2DS2+ NK cells from C*0102-IVDL vaccinated mice formed a discrete cluster to KIR2DS2+ NK cells from C*0102-AAA vaccinated mice, in contrast to KIR2DS2-NK cells (Fig. 2A). Those mice receiving C*0102-IVDL had upregulation of genes in pathways associated with cellular metabolism, as compared to those receiving control C*0102-AAA vaccination (Figure 2B). Differential gene expression analysis identified 54 differentially expressed genes (FDR<0.05) between the KIR2DS2+ NK cells from the C*0102-IVDL versus control groups (Table S1). These included upregulation in genes associated with: RNA binding and splicing; metabolism, especially glutathione metabolism; and regulation of IFN alpha (Fig. 2C). Additionally, we observed downregulation of genes related with histone H3 dimethylation at K4 (H3K4me2) and trimethylation at K27 (H3K27me3), consistent with a change in transcriptional regulation induced by vaccination (Fig. S4). H3K4Me2 epigenetic marks are associated with naïve T cells and H3K27me3 marks associated with reduced NK cell IFNγ secretion (32, 33). Thus downregulation of these pathways are consistent with activation or maturation of NK cells.

**Fig. 2:**
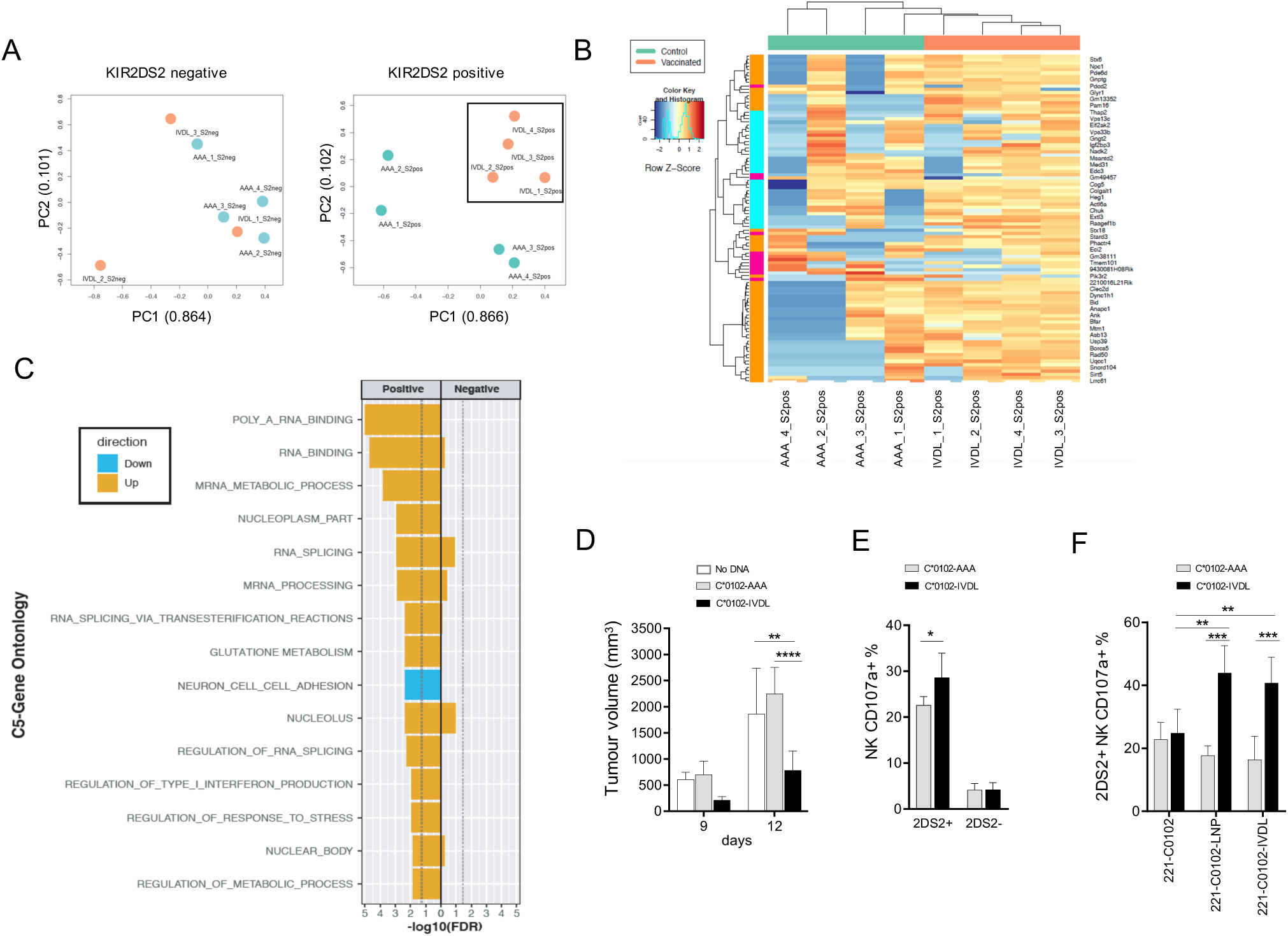
Preferential activation of KIR2DS2+ NK cells by peptide-based DNA vaccination. **A)** Principal component analysis (PCA) of whole NK cell transcriptomes from C*0102-IVDL and C*0102-AAA vaccinated mice. KIR2DS2+ NK cells from both groups are shown in the left panel and KIR2DS2-positive NK cells in the right panel. Counts were normalized and filtered using EdgeR. The first two components of the PCA are shown. **B)** Heatmap of the top 100 differentially expressed genes derived from the comparison of KIR2DS2 positive NK cells from C*0102-IVDL and C*0102-AAA vaccinated mice. **C)** EGSEA analysis of C5 signatures (GO) comparing KIR2DS2+ and KIR2DS2-NK cell populations in both C*0102-IVDL and control vaccinated mice. Effect significances were calculated individually for each arm of the study and the plot indicates the overall effects of vaccination on KIR2DS2+ NK cells in the C*0102-IVDL vaccinated mice (“positive”) as compared to the other three groups (“negative”). The color denotes the direction of the change and the size of the bar represents the -Log10(FDR). All categories shown were significant at FDR<0.05. **D)** KIR-Tg mice were injected subcutaneously with B16F10 melanoma cells on day 0 and then vaccinated intramuscularly with C*0102-IVDL (black bars) or C*0102-AAA (gray bars) on days 0 and 7 or untreated (white bars) and tumor volume measured. (n=4 mice per group: one of two independent experiments). **E**,**F)** Mice were injected intramuscularly with C*0102-IVDL (black bars) or C*0102-AAA (gray bars) on days 0 and day 7 and then NK cells purified from the spleens on day 14 for *in vitro* assays of activation. **E)** shows degranulation of KIR-Tg NK cells to B16F10 melanoma cells (n=4 mice per group). **F**) shows degranulation of KIR2DS2+ and KIR2DS2-KIR-Tg NK cells to human 721.221 cells expressing HLA-C*0102 alone (221-C*0102) or HLA-C*0102 in combination with the peptide: LNPSVAATL (221-C*0102-LNP) or IVDLMCHATF (221-C*0102-IVDL), (n=8 mice per group). Comparisons were by Students t-test (2 groups) or two-way ANOVA (more than 2 groups). For all plots *p < 0.05, **p< 0.01, ***p< 0.005, ****p< 0.001.

### A KIR targeting vaccine augments NK cell functions

To test for functional effects of our DNA vaccination strategy, KIR-Tg mice were inoculated subcutaneously in the flank with B16F10 melanoma cells and injected intramuscularly with DNA on the same day and one week later. Growth of B16F10 cells was significantly attenuated by day 12 in mice given C*0102-IVDL as compared to those given C*0102-AAA or unvaccinated (p<0.02) (Fig. 2D). In vitro, KIR2DS2+ NK cells, but not KIR2DS2-NK cells, from C*0102-IVDL-vaccinated mice had increased degranulation to B16F10 cells as compared to the control vaccinated mice (p<0.05) (Fig. 2E). As B16F10 cells do not express HLA-C these data indicate that DNA vaccination with C*0102-IVDL activates NK cells and induces MHC class I unrestricted responses.

To identify if peptide-specific NK cell responses were generated using this strategy, NK cells from vaccinated mice were tested against the MHC class I-negative target human cell line 721.221 transfected with either HLA-C or a construct of HLA-C in combination with the peptides LNPSVAATL and IVDLMCHATF (14). KIR2DS2+ NK cells from mice vaccinated with C*0102-IVDL demonstrated increased activity against 721.221 cells expressing HLA-C*0102 in combination with both KIR2DS2-binding peptides as compared to 721.221 cells transfected with HLA-C*0102 alone (p<0.01 for both LNP and IVDL targets) (Fig. 2F). No effect was observed for KIR2DS2-NK cells. Thus activation using a peptide:MHC strategy can generate both specific and cross-reactive peptide responses.

### KIR2DS2 recognizes a cancer associated antigen

Having identified that our strategy could activate NK cells to kill targets expressing a cross-reactive peptide, we investigated its potential as a therapeutic reagent. As KIR2DS2 has been associated with beneficial outcomes of cancer we sought to identify additional peptides that may be recognized by KIR2DS2. Hepatocellular carcinoma is one of the commonest cancers worldwide and is extremely difficult to treat pharmacologically (34). Activating KIR, including KIR2DS2, have been associated with a beneficial outcome to HCC (7). The Huh7 cell line is derived from human hepatocellular carcinoma (HCC) and has been used as a model in pre-clinical studies for HCC (35, 36). We used this as a model cell line to identify potential KIR2DS2-binding peptides. Huh7 cells express HLA-A*11, but HLA-B and HLA-C at undetectable levels on the cell surface and so, to provide an MHC class I ligand for KIR2DS2+ NK cells we transfected the cell line with HLA-C*0102 (37). We sequenced MHC class I bound peptides from Huh7 cells transfected with HLA-C*0102 (Huh7:C*0102) and from parental Huh7 cells following elution with the pan class I antibody W6.32. We identified ∼5800 and ∼7200 peptides from Huh7 and Huh7:C*0102 cells respectively (Table S2). The Huh7 immunopeptidome consisted mainly of peptides carrying a C-terminal lysine or arginine residue which is characteristic of the HLA-A*11 binding motif (Fig. 3A). In contrast, epitopes identified from the Huh7:C*0102 cell line contained two distinct populations, one with the HLA-A*11 binding motif and one with an HLA-C*0102 binding motif. From this comprehensive repertoire of HLA class I binders, only one peptide, NAPLVHATL, was identified that conformed to both the HLA-C*0102-binding motif (xxPxxxxxL) and the KIR2DS2-binding motif (xxxxxxATx). NAPLVHATL, which was identified by three independent, high-quality peptide-spectrum matches (Fig. 3B), is derived from the nuclear export protein exportin-1 (XPO1). This protein is upregulated in many hematological malignancies and solid tumors including HCC, is generally associated with a poor prognosis in cancer and has been successfully targeted in clinical trials of refractory multiple myeloma (38-43).

**Fig. 3.**
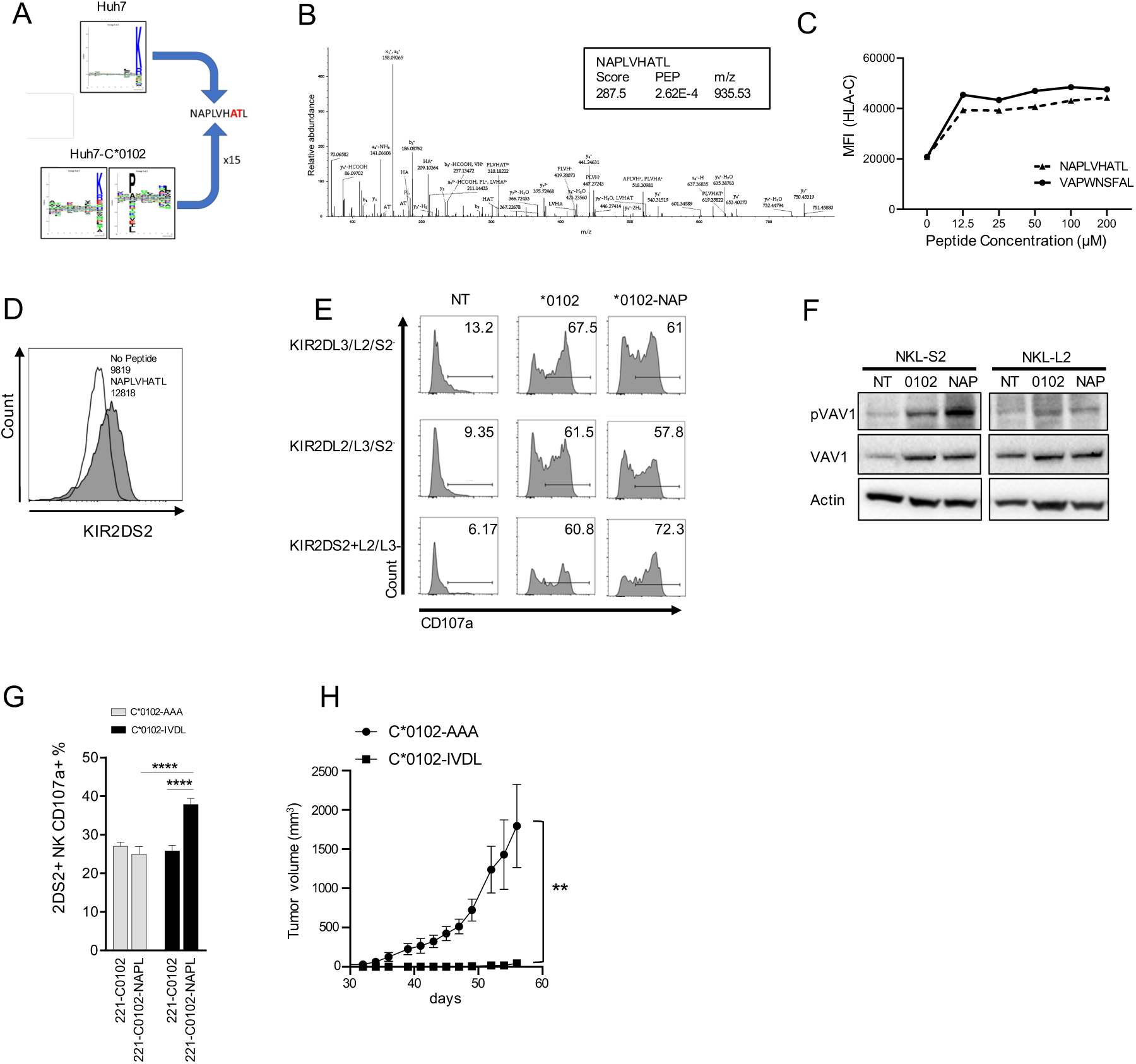
An XPO1 derived peptide is a ligand for KIR2DS2. **A)** SeqLogo plots of peptides eluted from Huh7 and Huh7:HLA-C*0102 cell lines. **B)** Representative and annotated ms2 spectrum that has been assigned to the peptide sequence NAPLVHATL. The resulting Byonic score, the posterior error probability (PEP) and the m/z value of the singly protonated species is shown in the inset. **C)** 721.174 cells were incubated with NAPLVHATL at the indicated concentrations, stained for HLA-C using the DT9 antibody and analyzed by flow cytometry. The mean fluorescence intensity (MFI) of DT9 staining compared to the control VAPWNSFAL peptide is shown. **D)** 721.174 were incubated with NAPLVHATL (100µM) and stained with a KIR2DS2-tetramer and analyzed by flow cytometry. Histogram plots of KIR2DS2-staining compared to a no peptide control are shown with MFIs indicated. **E)** 721.221 cells were transfected with HLA-C*0102 alone (C*0102) or in combination with the peptide NAPLVHATL (C*0102-NAP), and the cell lines used as targets for degranulation assays for IL-15 activated NK cells. CD107a expression on the indicated sub-populations of CD3-CD56+ NK cells was assessed by flow cytometry against these targets (NT, no target). For the gating strategy see Supplementary Figure 5. One representative flow cytometry plots of CD107a expression is shown from six donors tested. **F)** NKL-2DS2 or NKL-2DL2 cells were incubated with either no target (NT), 721.221:HLA-C*0102 (C*0102) or 721.221:HLA-C*0102+NAPLVHATL (NAP) cells for 5 minutes and assessed for VAV1 (Tyr174) phosphorylation by immunoblotting. A representative image from six experiments is shown. **G)** Mice were injected intramuscularly with C*0102-IVDL (black bars) or C*0102-AAA (gray bars on days 0 and day 7 and then NK cells purified from the spleens on day 14 for *in vitro* assays of activation against human target cell lines. Summarized degranulation data of KIR2DS2+ KIR-Tg NK cells to 721.221:HLA-C*0102+NAPLVHATL targets (n=4 mice per group) is shown. **H**) NK cells from KIR-Tg mice vaccinated either with C*0102-IVDL or C*0102-AAA as a peptide control, were adoptively transferred into NSG mice inoculated subcutaneously with Huh7:C*0102 hepatoma cells and tumor volume was measure (n=4 mice per group, one of two independent experiments). **p< 0.01 by ANOVA.

Data-independent acquisition mass spectrometry (DIA-MS) was used to quantify epitopes between the transfected Huh7:C*0102 and the parental Huh7 cell line (Table S3). The peptide NAPLVHATL was eluted at 15 fold greater levels in Huh7:C*0102 compared to Huh7 cells, consistent with it being presented by HLA-C*0102. Searching of the IEDB database revealed that this peptide has been previously eluted from C1R and 721.221 B-lymphoblastoid cell lines transfected with HLA-C*0102 (44, 45). In vitro, NAPLVHATL bound HLA-C*0102 expressed by TAP-deficient 721.174 cells to a similar level as VAPWNSFAL, a previously defined HLA-C*0102 binding peptide (46), and at saturating concentrations bound a KIR2DS2 tetramer (Fig. 3C and D, and Fig. S5).

KIR2DS2+/KIR2DL2/3-primary NK cells upregulated CD107a expression following incubation with 721.221 cells expressing NAPLVHATL in combination with HLA-C*0102 to a greater extent than other NK cell subsets (Fig. 3E and Fig. S6). The KIR-negative NKL cell line was transfected with KIR2DS2 (NKL-2DS2) or with KIR2DL2 (NKL-2DL2) and incubated with the 721.221 cells expressing HLA-C*0102 with or without NAPLVHATL. NKL-2DS2, but not NKL-2DL2 demonstrated enhanced VAV1 phosphorylation in the presence of NAPLVHATL, indicating that this peptide can induce activation of NKL cells in a KIR2DS2 specific manner (Figure 3F).

To test if our vaccination strategy induced specific responses against NAPLVHATL we tested the activity of NK cells from C*0102-IVDL-vaccinated KIR-Tg mice against transfected 721.221 cells. This demonstrated increased degranulation of KIR2DS2+ NK cells against 721.221 cells expressing NAPLVHATL in combination with HLA-C*0102 as compared to those expressing HLA-C*0102 only (p<0.05) (Figure 3G). Thus, NAPLVHATL is recognized by KIR2DS2 in the context of HLA-C*0102, and this interaction activates NK cells. To determine if this response translated into a functional benefit we performed an adoptive transfer study. KIR-Tg mice were vaccinated with two doses of the DNA vaccine C*0102-IVDL or the control vaccine weekly, and then purified splenic NK cells containing <1% CD3+ T cells, were adoptively transferred into immunodeficient NSG mice, which had been inoculated with Huh7-C*0102 cells. We observed a significantly delayed growth of the tumor in mice that received C*0102-IVDL-stimulated NK cells compared to control vaccine concomitant (Fig. 3H). Thus stimulation of NK cells via KIR2DS2 can generate anti-cancer reactivity against HLA-C expressing human tumor cells. To test if endogenously expressed XPO1 was recognized by KIR2DS2, XPO1 was knocked down in Huh7:C*0102 cells using siRNA (Fig. 4A). NKL-2DS2 cells, but not NKL-2DL2, killed XPO1 knockdown Huh7:C*0102 cells less efficiently than those transfected with control siRNA at all effector:target (E:T) ratios tested (p<0.001) (Fig. 4B and C). Furthermore XPO1 knockdown decreased CD107a expression on KIR2DS2+ primary NK cells as compared to KIR2DS2-NK cells (Fig. 4D and E). Thus, endogenously expressed XPO1 is recognized specifically by KIR2DS2+ NK cells.

**Fig. 4.**
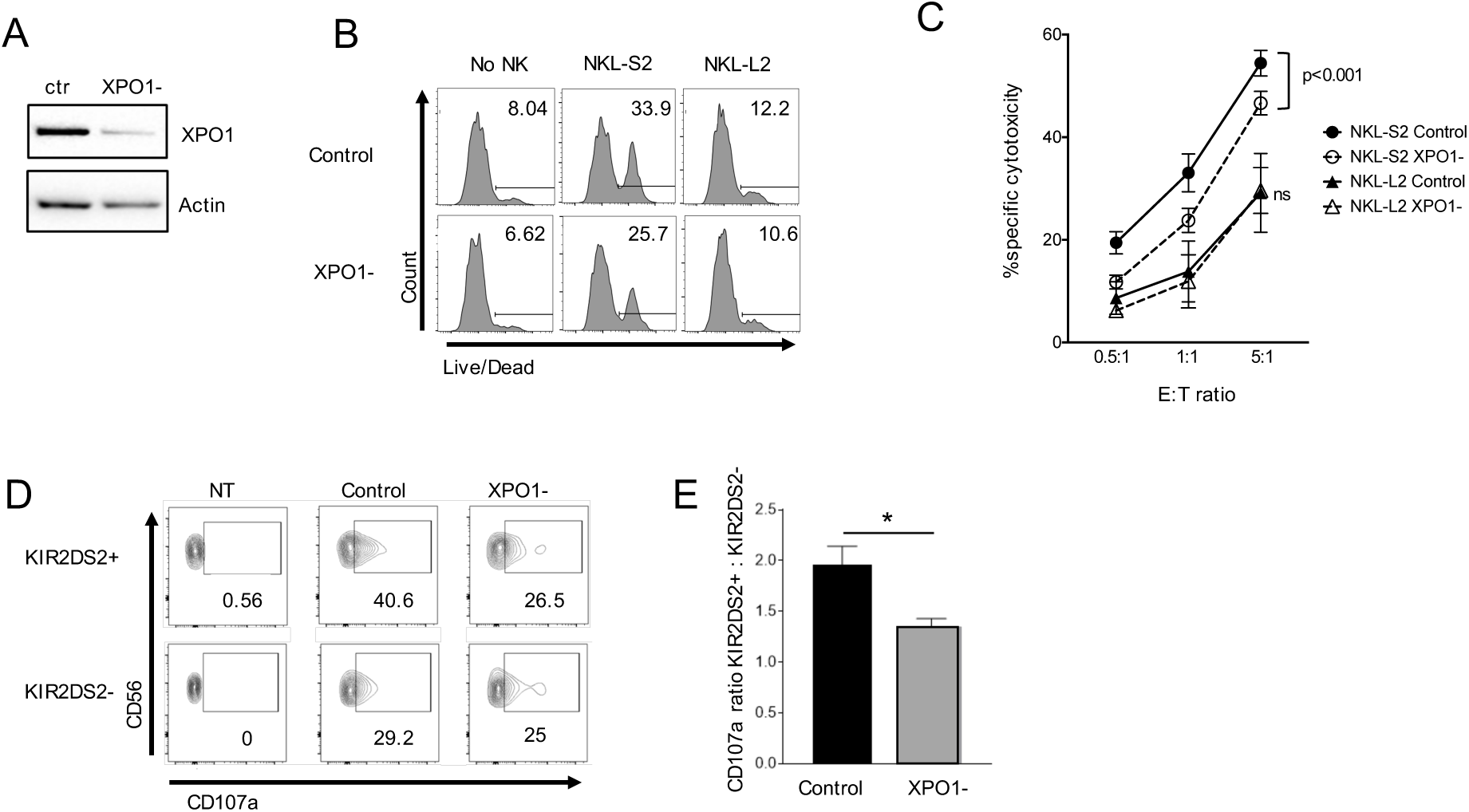
Silencing of XPO1 expression in Huh7:HLA-C*0102 cells reduces KIR2DS2+ but not KIR2DS2-NK cell activation. **A)** Representative immunoblot of XPO1 silencing following XPO1-targeting siRNA treatment of Huh7:HLA-C*0102 cells. **B)** and **C)** Huh7:HLA-C*0102 cells were treated with control or XPO1 targeting siRNA and used as targets in cytotoxicity assays with NKL-S2 or NKL-L2 cells. Killing was determined using the LIVE/DEAD™ stain. Representative plots at an effector:target ratio of 5:1 are shown in **B)** and the mean specific cytotoxicity and SEM from five independent experiments are shown **C)**, (***p<0.005)**. D)** and **E)** Huh7:HLA-C*0102 cells were treated with control or XPO1-targeting siRNA and used as targets for KIR2DS2+ NK cells in CD107a degranulation assays. CD107a expression in KIR2DS2+ or KIR2DS2-CD3-CD56+ NK cells determined using the 1F12 antibody in KIR2DL2/S2 homozygous donors was analyzed by flow cytometry. Representative contour plots are shown in **D)** and the ratio of CD107a positivity in KIR2DS2+ to KIR2DS2-NK cells from three donors is shown **E)** (*p<0.05).

## Discussion

We describe proof-of-concept evidence that NK cells can be activated through peptide:MHC to generate functional immune responses. In our model, KIR2DS2-mediated activation generated a cytotoxic immune response against targets both with and without a cognate KIR2DS2-ligand. However we did not observe specific proliferation of KIR2DS2+ NK cells and the levels of KLRG1 expression are lower than would be found during a viral infection such as MCMV, which is not surprising as the same cytokine milieu may not be created by DNA vaccination (47). However, it is consistent with co-expression of the antigen-specific receptor Ly49H and KLRG1 noted following MCMV infection (48). This strategy may have implications for cancer therapy. In particular we provide evidence that tumors may be directly susceptible to KIR2DS2-mediated killing by identifying a human cancer-associated peptide from XPO1 that is an HLA-C restricted ligand for KIR2DS2. XPO1 (exportin-1/CRM1) is thought to function as the sole nuclear exporter for many different tumor suppressor and growth regulatory proteins including p53, p27, FOXO1, IkB, cyclin B1, cyclin D1 and survivin (38). It is overexpressed and associated with poor prognosis in multiple cancers including hematological malignancies and difficult to treat cancers such as pancreatic cancer and HCC (41, 43, 49). Inhibition of XPO1 is gaining traction as a cancer therapeutic target following clinical trials, and our data suggest it could also be a target for KIR2DS2+ NK cells (40). Previous work has also shown that KIR2DS2+ NK cells can recognize several different cancer targets *in vitro*, including cell lines derived from prostate, breast and ovarian carcinomas (27). However this recognition was not specific to KIR2DS2 and was beta-2-microglobulin independent, suggesting that there may also be non-peptide:MHC class I ligands for KIR2DS2. Additionally, KIR2DS2+ NK cells appear to have a greater potential to mediate ADCC *in vitro* and *in vivo* (28, 29), therefore KIR2DS2 may be an attractive target for cancer immunotherapy in combination with antibody based therapeutics.

The *in vivo* studies demonstrated that KIR2DS2 could be targeted using a peptide:MHC specific approach. We observed activation of both KIR2DS2+ and KIR2DS2-NK cells *in vivo*, but activation was preferential for KIR2DS2+ NK cells and, *in vitro*, peptide specific recognition of 721.221 targets was only noted for splenocytes from mice vaccinated with IVDLMCHATF. Our data is consistent with a model of activation through one receptor and killing mediated by a different receptor; for instance killing of the B16 melanoma cell line is mediated by NKp46, and hence not MHC class I restricted (50). Additionally, we demonstrate peptide cross reactivity of KIR2DS2, as mice vaccinated with IVDLMCHATF were able to recognize the XPO1 peptide NAPLVHATL, and also the HCV peptide LNPSVAATL, consistent with KIR2DS2 recognizing peptides with an AT motif at the carboxy-terminal −1 and −2 positions. Thus, in order to take our findings to the clinic it is not necessary to identify the ligand on the cancerous cell, or to match for HLA class I as KIR2DS2 may bind other group 1 HLA class I allotypes such as HLA-C*0304 (14, 16). However, it may be necessary to refine the strategy by the addition of cytokines such as IL-15 to induce proliferation of KIR2DS2-positive NK cells. The broad specificity of KIR2DS2 also provides an advantage for peptide-based NK cell therapy over T cell pMHC therapeutics which require more precise pMHC matching. Additionally as the strategy provides both the peptide and the MHC class I ligand, then no HLA class I matching is required. As NK cells are involved in the early immune response to viral infections, and KIR2DS2 recognizes peptides derived from many different viruses, then targeting KIR2DS2 by DNA vaccination may also form part of an anti-viral therapeutic strategy to reduce infection and transmission in the early stages of infection. In conclusion, our work identifies a novel cancer associated ligand for KIR2DS2 and provides evidence that KIR2DS2 ligation inhibits tumor growth in vivo. This data provides a rationale for targeting activating KIR as a novel NK cell based therapy for human cancer.

## Materials and Methods

### Mice vaccination and tumor models

KIR transgenic mouse expressing a complete human KIR B haplotype, on a C57BL6 background MHC class I–deficient Kb^-/-^ Db^-/-^ were a kind gift from J Van Bergen and kept under specific pathogen-free conditions (31). DNA constructs were made expressing HLA-C*0102 alone or linked with the T2A self-cleaving peptide, and peptide and cloned into pIB2 (14). For the B16 model, mice were injected intramuscularly with 50µg DNA. 2.5×10^5^ B16F10 cells into the mice left flank, and 50ug of DNA plasmid on days 0 and 7. For the Huh7 model, NSG mice were injected with 2×10^6^ Huh7:C*0102 cells subcutaneously and then10^6^ purified NK cells from 2 weeks vaccinated KIR-Tg mice spleen were injected intravenously on days 0 and 14. NK cells were purified using MACS technology (Miltenyi Biotec, NK cell isolation kit II). Purity was > 90% NK cells with < 3% CD3^+^ T cells.

### Cell lines

HLA class I-deficient 721.221 lymphoblastoid EBV-B cells were cultured in R10 medium (RPMI 1640 supplemented with 1% penicillin-streptomycin (Life Technologies) and 10% heat inactivated fetal bovine serum (FBS; Sigma). 721.221 cells were transduced with the pIB2 constructs to express HLA-C*0102 alone or together with peptides LNPSVAATL, IVDLMCHATF or NAPLVHATL. For tetramer staining 721.174 cells were cultures in R10 incubated overnight with peptides and then stained KIR2DS2 tetramers as previously described (14). B16F10 cells were cultured in Dulbecco’s modified Eagle’s medium (DMEM) with 1% penicillin-streptomycin (Life Technologies) and 10% FBS.

### Flow cytometry analyses

Murine splenocytes and intra-hepatic lymphocytes were stained using anti-mouse CD3ε-PE, NK1.1-BV421, CD11b-APC-Cy5, CD27-BV510, KLRG1-PECy7, (Biolegend). The 1F12-FITC antibody was used to selectively stain KIR2DS2 (51). For CD107a assays, freshly isolated splenocytes were cultured with anti-CD107a AF647 (Biolegend) and GolgiStop™ (BD Biosciences) prior to staining. Cells were stained with 1F12-FITC, CD3-PE, NK1.1-BV421 (Biolegend). For the IFN-γ secretion assay, splenocytes were surface stained with 1F12-FITC, CD3-PE, NK1.1-BV421 antibodies, fixed and permeabilized using BD Cytofix/Cytoperm buffers and then stained with anti-mouse IFNγ-APC (Biolegend). Events were acquired on Aria II (BD Biosciences) using the FACSDiva software (BD Biosciences) and analyzed with FlowJo software (Treestar).

Human PBMC were obtained with full ethical approval from the National Research Ethics Committee reference 06/Q1701/120. IL-15 stimulated PBMCs were incubated with 721.221 or Huh7:HLA-C*0102 cells for 4 hours, anti-CD107a-AF647 (Biolegend), GolgiStop added and cells analyzed by flow cytometry. For Huh7:HLA-C*0102 cytotoxicity assays target cells were co-incubated with the indicated NK cell population for 4 hours. Cells were then stained with LIVE/DEAD™ stain (ThermoFisher Scientific) and analyzed by flow cytometry, gating on the target cell population identifiable by GFP within the HLA-C*0102 construct. For XPO1 knockdown, Huh7:HLA-C*0102 cells were transfected using Jetprime (Polyplus, UK) reagent with siRNA control or siRNA targeting XP01 (ThermoFisher Scientific). XP01 expression was analyzed by immunoblotting after 48 hours and cells were used in cytotoxicity and degranulation assays 48 hours post transfection.

### Immunoblotting

NKL-KIR2DS2 or NKL-KIR2DL2 were co-incubated with 721.221:HLA-C*0102 or 721.221:HLA-C*0102+NAPLVHATL cells as indicated for 5min at an E:T ratio of 1:1. Cells were then lysed in NP40 Cell Lysis Buffer (Fisher Scientific UK Ltd) and analyzed by immunoblotting. Antibodies recognizing phospho-VAV1, VAV1 and actin (ThermoFisher Scientific) were used with HRP-conjugated antibodies and visualized using ChemiDoc-It Imaging system (BioRad). Bands were quantified using ImageJ software.

### RNA Seq data analysis and processing

RNA was isolated using the RNeasy Kit (QIAGEN) and prepared the QIAseq UPX 3’ Transcriptome Kit (QIAGEN). 10ng purified RNA was used for the NGS libraries. The library pool was sequenced on a NextSeq500 instrument. Raw data was de-multiplexed and FASTQ files generated using bcl2fastq software (Illumina inc.). FASTQ data were checked using the FastQC tool (https://www.bioinformatics.babraham.ac.uk/projects/fastqc/). For differential gene expression analysis, raw counts from RNA-Seq were processed in Bioconductor package EdgeR(52), variance was estimated and size factor normalized using TMM. Genes with minimum 4 reads at minimum 40% samples were included in the downstream analyses. Differentially expressed genes (DEG) were identified applying significance threshold FDR p<0.05. Blind, normalized log2 values calculated by EdgeR were used for principal component analysis and to calculate Euclidean distances for hierarchical clustering using Ward’s method. For heatmaps the normalized log2 values of all high-fold change peaks were used to hierarchically cluster peak regions into seven clusters, with the top 100 most variable genes (based on calculated variance across all samples). Gene ontology and pathway enrichment analysis were done using CAMERA (53), Ensemble of Gene Set Enrichment Analyses (EGSEA) (54) and ToppGene (55). All analyses used default settings considering mouse orthologs from the MSigDB v5.2 databases retrieved from http://bioinf.wehi.edu.au/software/MSigDB/. Only pathway terms with a minimum of 25 genes were considered and used for multiple-hypothesis correction. Enriched pathways were filtered for those that showed Padj < 0.05 for both P values as calculated for the contrast C*0102-IVDL vs C*0102-AAA in KIR2DS+ and KIR2DS2-NK cell populations. Pathway results were further filtered on those that showed the lowest p.adj values.

### Peptide elution, identification and quantification

HLA class I peptides were eluted from Huh7 and Huh7:HLA-C*0102 cells as previously described before (56-58). HLA-peptide eluates were loaded onto C18 RP-HPLC column (Chromolith Speed Rod; Merck). The bound peptides were separated using increasing concentration of 80% acetonitrile and 0.1% trifluoroacetic acid (80% ACN / 0.1% FA). Peptide-containing fractions were collected, and reconstituted with 0.1% formic acid (FA). Indexed retention time (iRT) peptides (Escher et al., 2012) were spiked in for retention time alignment. Peptides were then loaded onto a Dionex UltiMate 3000 RSLCnano system via an Acclaim PepMap 100 trap column onto an Acclaim PepMap RSLC analytical column (ThermoFisher Scientific), separated using increasing concentrations of 80% ACN/0.1% FA and analyzed with a QExactive mass spectrometer (ThermoFisher Scientific). HLA-bound peptides were quantified using data-independent acquisition (DIA) on an Orbitrap Fusion Tribrid mass spectrometer (ThermoFisher Scientific) coupled to an identical LC setup. 50 sequential DIA windows with an isolation width of 12 m/z between 375 - 975 m/z were acquired in 2 consecutive injections following a full ms1 scan (resolution: 120.000; AGC target: 4e5; maximum IT: 50 ms; scan range: 375-1575 m/z). Acquired DDA .raw files were searched against the human UniProtKB/SwissProt database (v2012_07) using Byonic (Protein Metrics) embedded in Proteome Discoverer (ThermoFisher Scientific) to obtain peptide sequence information. Only peptides identified at a false discovery rate (FDR) of 1% based on a decoy database were considered for further analysis. Spectronaut Orion (Biognosys) was used to create the corresponding spectral library as well as to evaluate all DIA data using in-house, peptide-centric parameters.

### Statistical analysis

Experimental statistical analyses were performed using Graph-Pad Prism 7.0 software. Student two-tailed t test was used for comparison between two groups and two-way ANOVA with post-hoc analysis were used to compare more than two groups. Data were considered statistically significant at p< 0.05. For the tumor model statistical comparisons between survival to the humane end point was performed by Log-rank test (Mantel-cox).

## Acknowledgements

We would like to than Jeroen van Bergen and John Trowsdale for KIR-Tg mice and Mohammed Naiyer for technical assistance.

---

**Figure S1:**
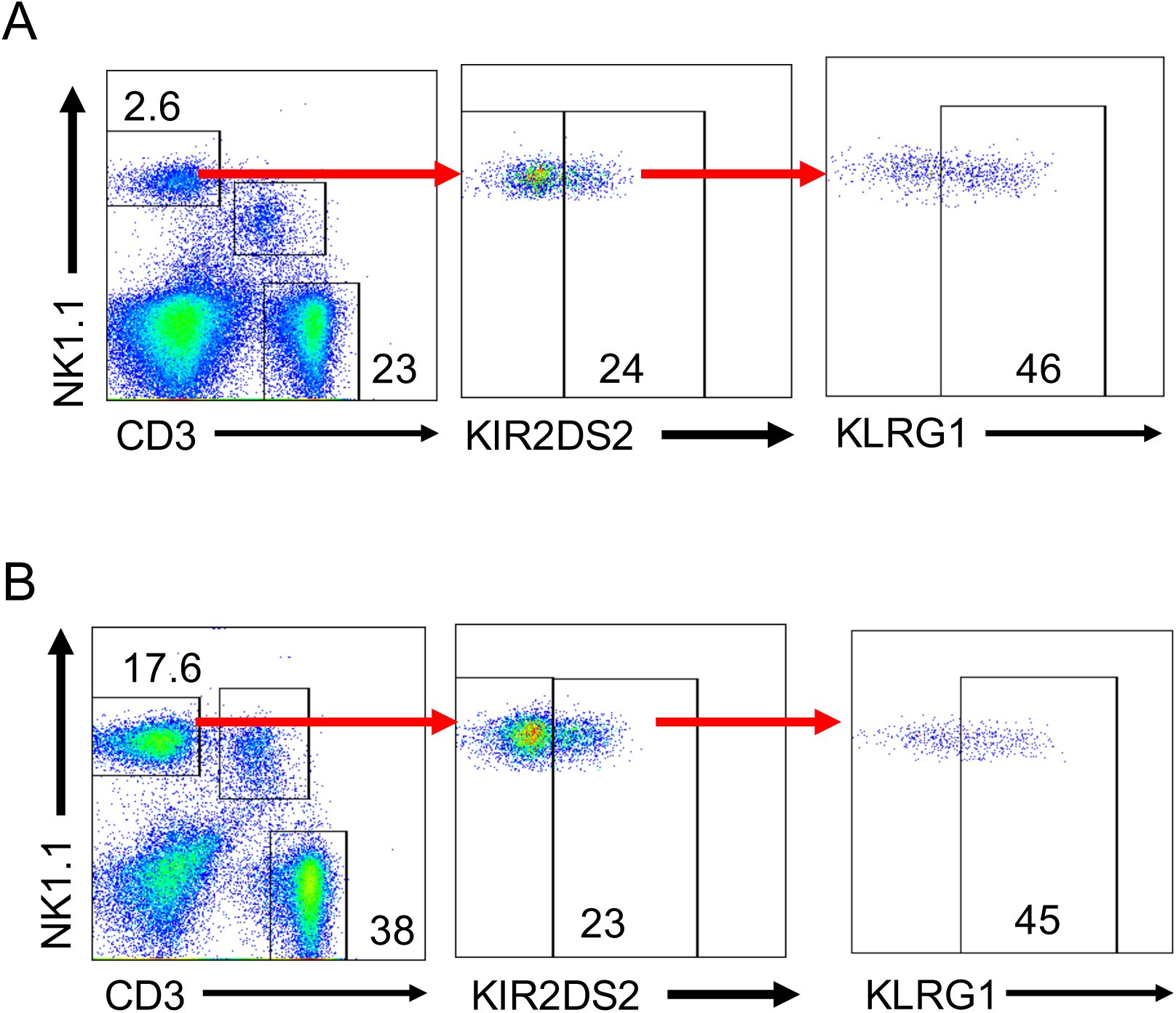
Gating strategy for KLRG1 on KIR2DS2-positive NK cells. Gating strategy for KLRG1 on KIR2DS2+ NK cells derived from spleens (**A**) and livers (**B**) of KIR-Tg mice. KIR2DS2+ NK cells were identified using the antibody 1F12.

**Figure S2:**
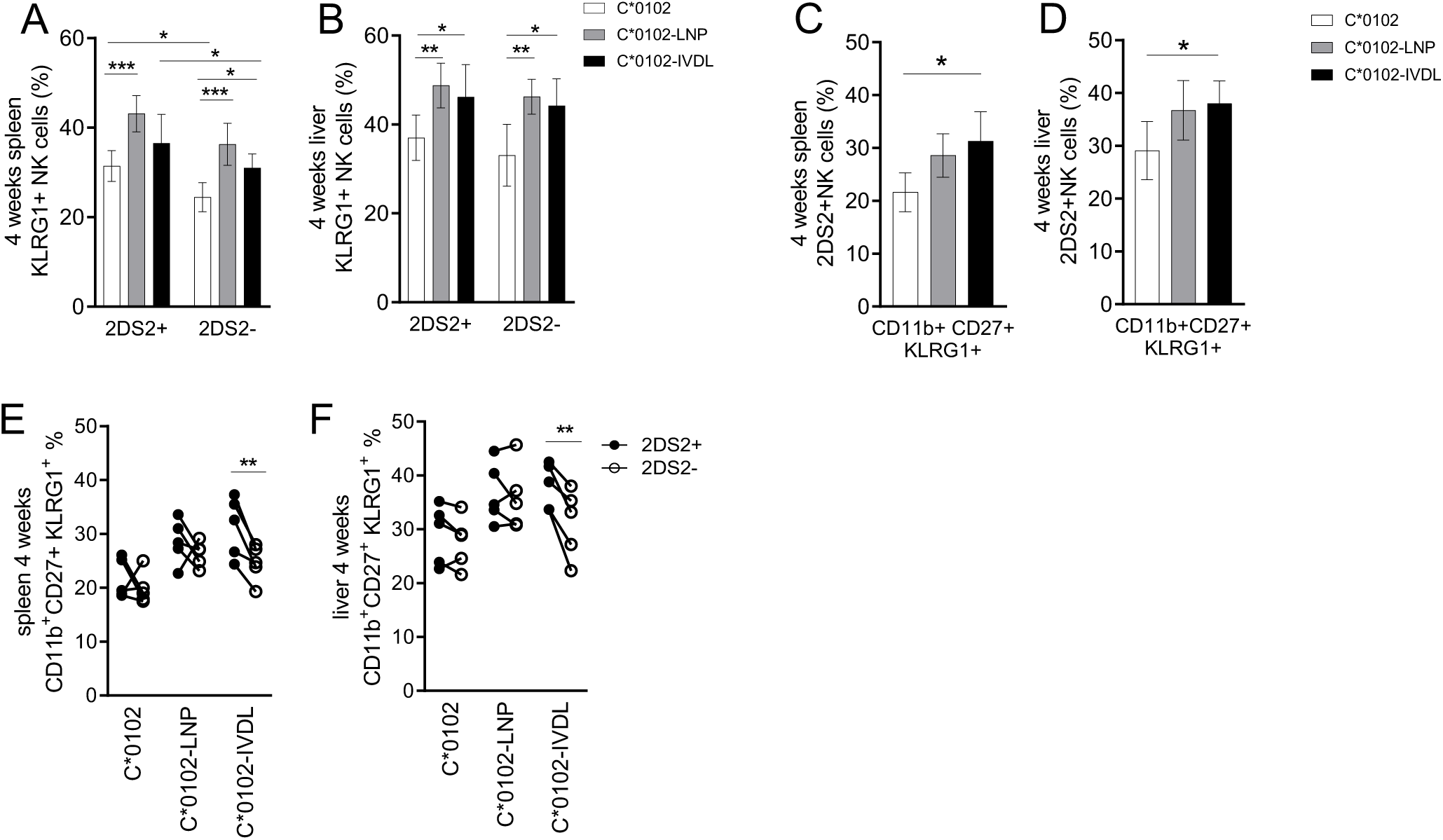
Activation of NK cells after DNA vaccination for 4 weeks. KIR-Tg mice were injected intramuscularly weekly for 4 weeks with DNA constructs encoding HLA-C*0102 (white bars), HLA-C*0102-LNPSVAATL (C*0102-LNP; gray bars) or HLA-C*0102-IVDLMCHATF (HLA-C*0102-IVDL; black bar). KLRG1 expression was measured on KIR2DS2+ NK cells and CD11b, CD27 subsets. **A, B**) KLRG1 frequencies on KIR2DS2+ and KIR2DS2-NK cells in spleen **(A)** and livers **(B). C, D).** Frequency of KLRG1 expression on CD11b+CD27+ NK cells in the KIR2DS2+ NK cell subpopulations in the spleens (**C)** and livers **(D)** of vaccinated KIR-Tg mice. **E, F)** Comparison of KLRG1+ expression on CD11b+CD27+ within the KIR2DS2+ and KIR2DS2-NK cells in the spleens (**E**) and livers (**F**) of vaccinated mice. For all experiments n=4 mice per group. A paired t test was used for comparisons between two groups and a 2-way ANOVA with correction used when comparing more than two groups. For all plots *p< 0.05, **p< 0.01, ***p< 0.005.

**Figure S3:**
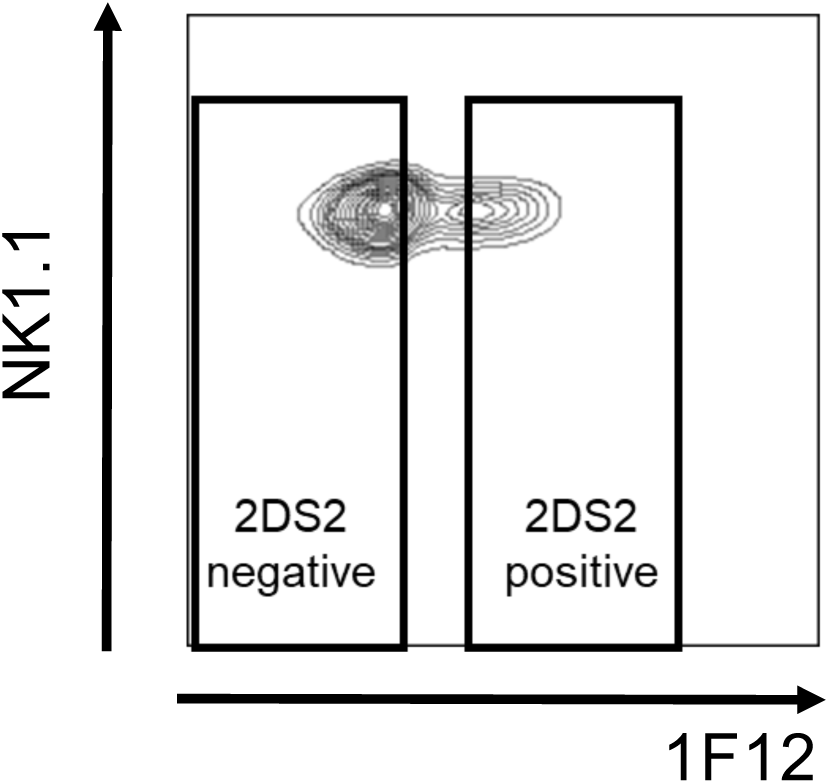
Gating strategy for sorting KIR2DS2-positive and -negative NK cells for analysis by RNA seq. Splenocytes from C*0102:IVDL and C*0102:AAA vaccinated mice were isolated and sorted by flow cytometry for analysis by RNAseq. Shown is a flow cytometry plot of the gating strategy used to delineate KIR2DS2+ from KIR2DS2-NK cells. Cells are gated on the CD3-, NK1.1+ sub-population.

**Figure S4:**
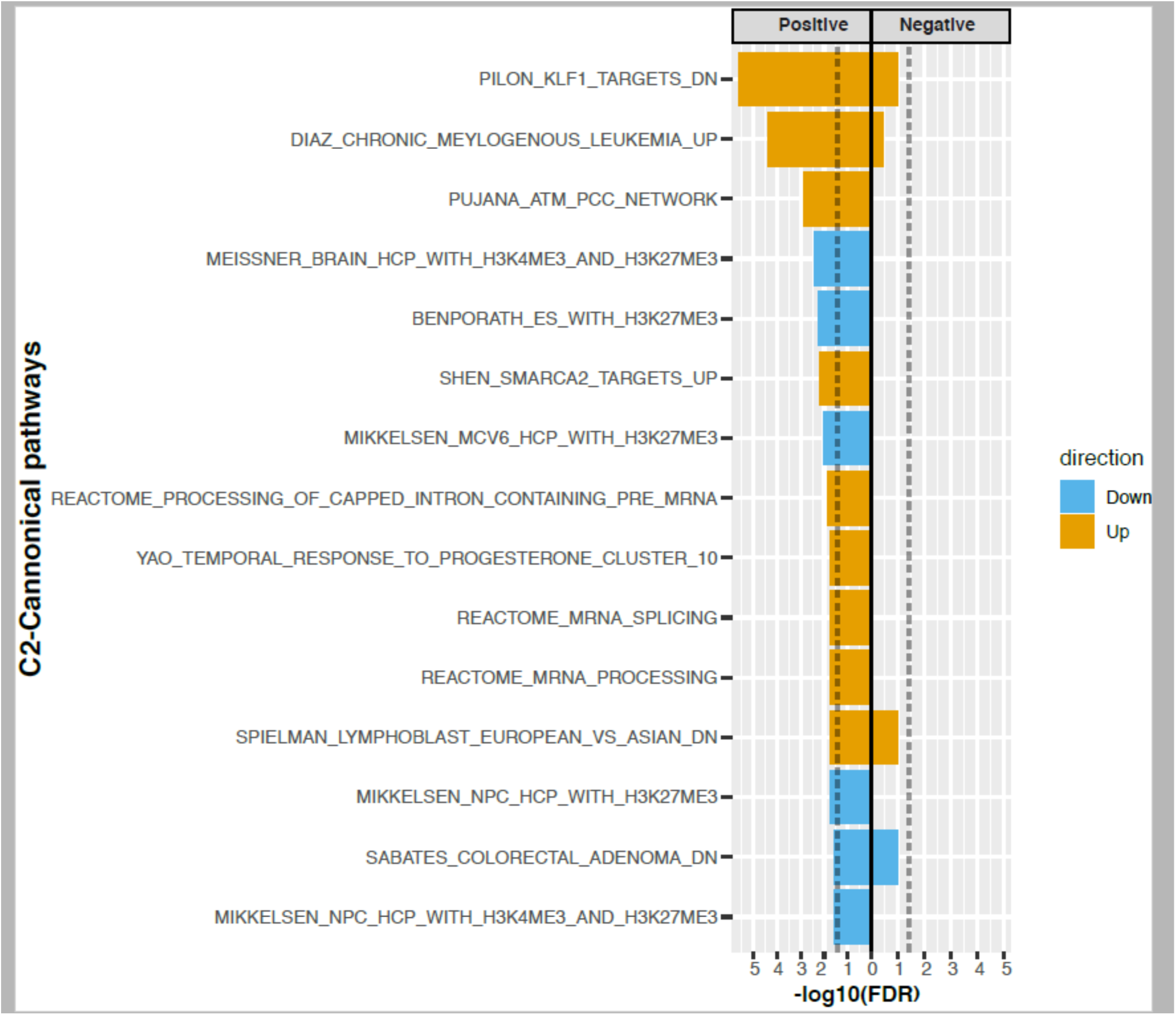
EGSEA analysis of differentially expressed genes comparing 2DS2-positive NK cells from C*0102-IVDL versus C*0102-AAA vaccinated mice. KIR2DS2+ and KIR2DS2-NK from mice vaccinated with C*0102-IVDL or the control vaccine C*0102-AAA were isolated by flow cytometry and analysed by RNAseq. Shown is an EGSEA analysis of the differentially expressed genes (FDR<0.05) between the 2DS2-positive NK cells from the C*0102-IVDL versus the 2DS2-positive NK cells from the C*0102-AAA control group, indicating upregulation of the H3K4me2 and H3K27me3 pathways.

**Figure S5:**
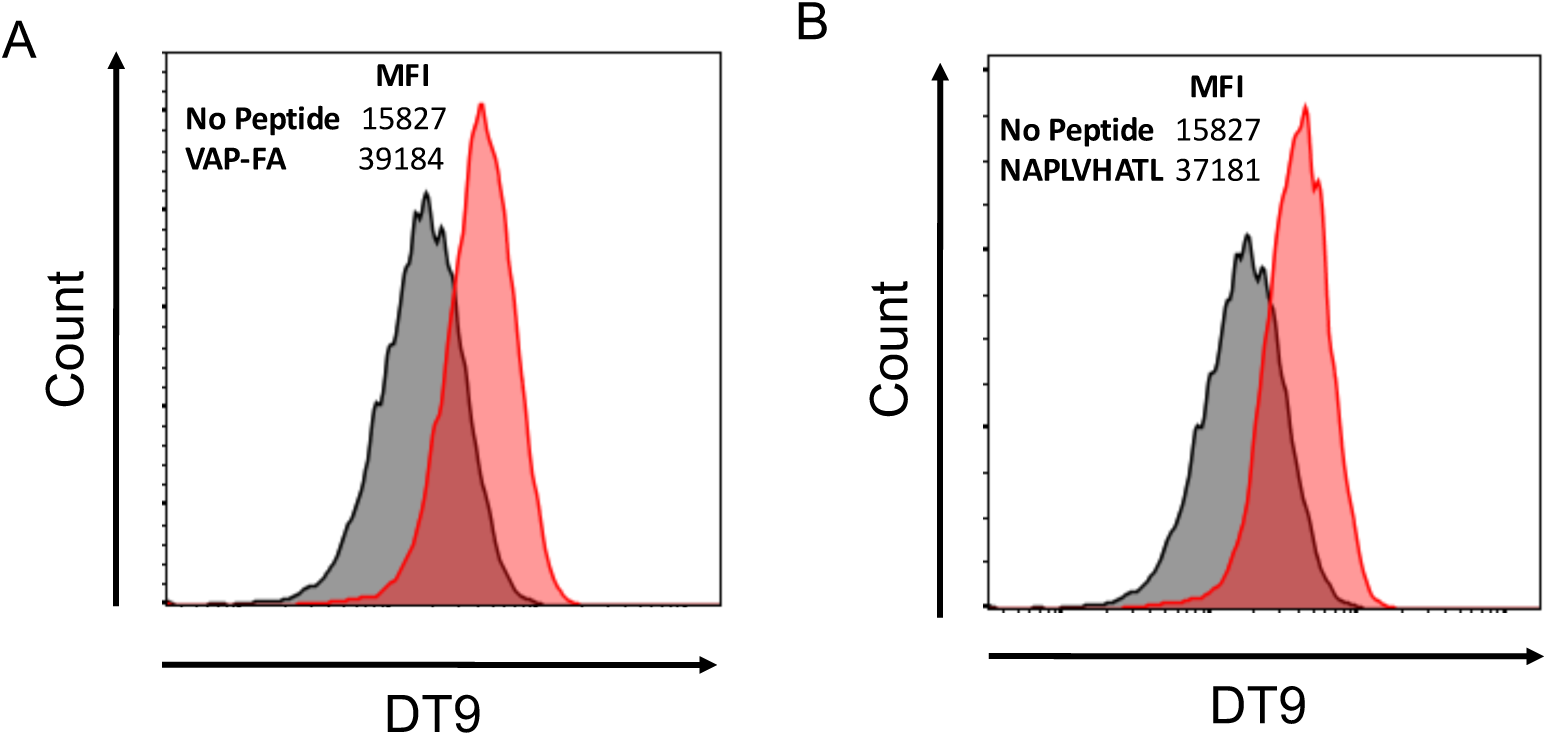
Flow cytometry plot HLA-C stabilisation by NAPLVHATL. Representative FACs plots of HLA-C staining of 721.174 cells either with no peptide (black) or loaded with 200µM of A) VAP-FA peptide (red) or B) NAPLVHATL peptide (red).

**Figure S6:**
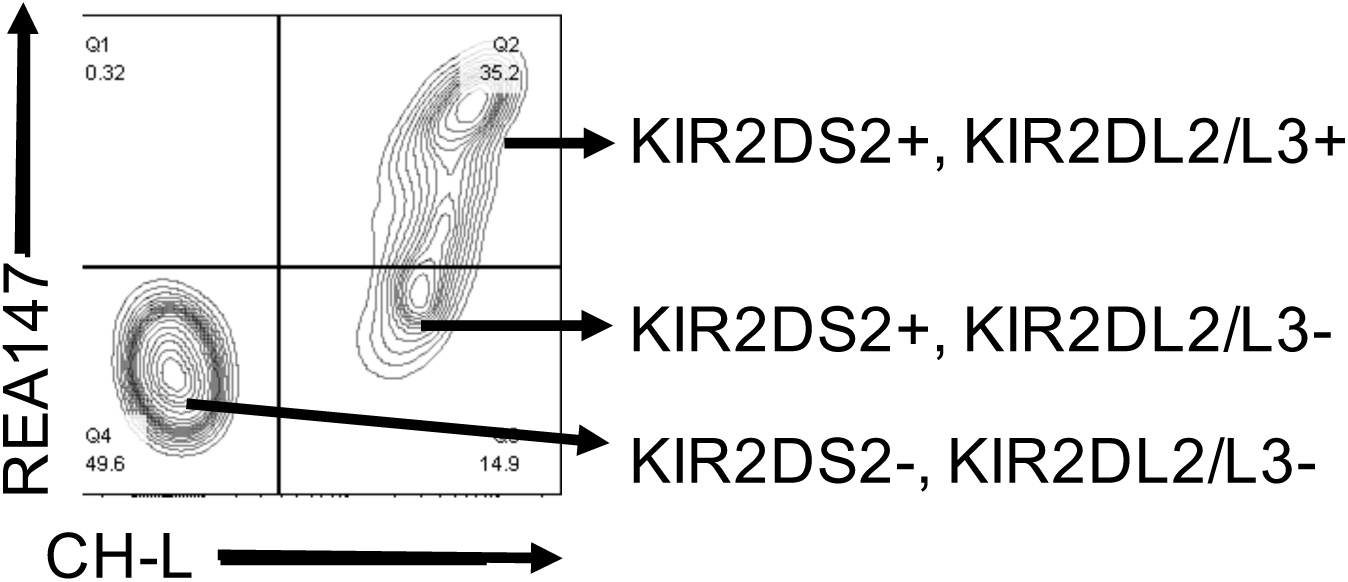
KIR2DS2 gating strategy for human PBMCs. Gating strategy to identify KIR2DS2+KIR2DL2/3-NK cells using the antibodies CH-L and REA147 as defined by Blunt et al (*57*). KIR2DS2+KIR2DL2/3-NK cells are CH-L positive and REA147 negative. Cells are gated on CD3-CD56+ lymphocytes

## References

1. Demaria O, et al. (2019) Harnessing innate immunity in cancer therapy. Nature 574(7776):45–56.

2. Cooley S, Parham P, & Miller JS (2018) Strategies to activate NK cells to prevent relapse and induce remission following hematopoietic stem cell transplantation. Blood 131(10):1053–1062.

3. Hu W, Wang G, Huang D, Sui M, & Xu Y (2019) Cancer Immunotherapy Based on Natural Killer Cells: Current Progress and New Opportunities. Front Immunol 10:1205.

4. Vilches C & Parham P (2002) KIR: diverse, rapidly evolving receptors of innate and adaptive immunity. Annu Rev Immunol 20:217–251.

5. Leone P, De Re V, Vacca A, Dammacco F, & Racanelli V (2017) Cancer treatment and the KIR-HLA system: an overview. Clin Exp Med 17(4):419–429.

6. Bao X, et al. (2018) HLA and KIR Associations of Cervical Neoplasia. J Infect Dis 218(12):2006–2015.

7. Cariani E, et al. (2013) HLA and killer immunoglobulin-like receptor genes as outcome predictors of hepatitis C virus-related hepatocellular carcinoma. Clin Cancer Res 19(19):5465–5473.

8. Carrington M, et al. (2005) Hierarchy of resistance to cervical neoplasia mediated by combinations of killer immunoglobulin-like receptor and human leukocyte antigen loci. J Exp Med 201(7):1069–1075.

9. Cornillet M, et al. (2019) Imbalance of Genes Encoding Natural Killer Immunoglobulin-Like Receptors and Human Leukocyte Antigen in Patients With Biliary Cancer. Gastroenterology 157(4):1067–1080 e1069.

10. Impola U, et al. (2014) Donor Haplotype B of NK KIR Receptor Reduces the Relapse Risk in HLA-Identical Sibling Hematopoietic Stem Cell Transplantation of AML Patients. Front Immunol 5:405.

11. Almalte Z, et al. (2011) Novel associations between activating killer-cell immunoglobulin-like receptor genes and childhood leukemia. Blood 118(5):1323–1328.

12. Vey N, et al. (2018) A phase 1 study of lirilumab (antibody against killer immunoglobulin-like receptor antibody KIR2D; IPH2102) in patients with solid tumors and hematologic malignancies. Oncotarget 9(25):17675–17688.

13. Stewart CA, et al. (2005) Recognition of peptide-MHC class I complexes by activating killer immunoglobulin-like receptors. Proc Natl Acad Sci U S A 102(37):13224–13229.

14. Naiyer MM, et al. (2017) KIR2DS2 recognizes conserved peptides derived from viral helicases in the context of HLA-C. Sci Immunol 2(15).

15. Sim MJW, et al. (2019) Human NK cell receptor KIR2DS4 detects a conserved bacterial epitope presented by HLA-C. Proc Natl Acad Sci U S A 116(26):12964–12973.

16. David G, et al. (2013) Large spectrum of HLA-C recognition by killer Ig-like receptor (KIR)2DL2 and KIR2DL3 and restricted C1 SPECIFICITY of KIR2DS2: dominant impact of KIR2DL2/KIR2DS2 on KIR2D NK cell repertoire formation. J Immunol 191(9):4778–4788.

17. Boyington JC, Brooks AG, & Sun PD (2001) Structure of killer cell immunoglobulin-like receptors and their recognition of the class I MHC molecules. Immunol Rev 181:66–78.

18. Das J & Khakoo SI (2015) NK cells: tuned by peptide? Immunol Rev 267(1):214–227.

19. Blunt MD, et al. (2019) A novel antibody combination to identify KIR2DS2(high) natural killer cells in KIR2DL3/L2/S2 heterozygous donors. HLA 93(1):32–35.

20. Cooley S, et al. (2009) Donors with group B KIR haplotypes improve relapse-free survival after unrelated hematopoietic cell transplantation for acute myelogenous leukemia. Blood 113(3):726–732.

21. Cooley S, et al. (2014) Donor killer cell Ig-like receptor B haplotypes, recipient HLA-C1, and HLA-C mismatch enhance the clinical benefit of unrelated transplantation for acute myelogenous leukemia. J Immunol 192(10):4592–4600.

22. Bachanova V, et al. (2016) Donor KIR B Genotype Improves Progression-Free Survival of Non-Hodgkin Lymphoma Patients Receiving Unrelated Donor Transplantation. Biol Blood Marrow Transplant 22(9):1602–1607.

23. Sekine T, et al. (2016) Specific combinations of donor and recipient KIR-HLA genotypes predict for large differences in outcome after cord blood transplantation. Blood 128(2):297–312.

24. Alomar SY, et al. (2017) Association of the genetic diversity of killer cell immunoglobulin-like receptor genes and HLA-C ligand in Saudi women with breast cancer. Immunogenetics 69(2):69–76.

25. Beksac K, Beksac M, Dalva K, Karaagaoglu E, & Tirnaksiz MB (2015) Impact of “Killer Immunoglobulin-Like Receptor /Ligand” Genotypes on Outcome following Surgery among Patients with Colorectal Cancer: Activating KIRs Are Associated with Long-Term Disease Free Survival. PLoS One 10(7):e0132526.

26. Wisniewski A, et al. (2012) KIR2DL2/S2 and HLA-C C1C1 genotype is associated with better response to treatment and prolonged survival of patients with non-small cell lung cancer in a Polish Caucasian population. Hum Immunol 73(9):927–931.

27. Thiruchelvam-Kyle L, et al. (2017) The Activating Human NK Cell Receptor KIR2DS2 Recognizes a beta2-Microglobulin-Independent Ligand on Cancer Cells. J Immunol 198(7):2556–2567.

28. Gras Navarro A, et al. (2014) NK cells with KIR2DS2 immunogenotype have a functional activation advantage to efficiently kill glioblastoma and prolong animal survival. J Immunol 193(12):6192–6206.

29. Siebert N, et al. (2016) Neuroblastoma patients with high-affinity FCGR2A, −3A and stimulatory KIR 2DS2 treated by long-term infusion of anti-GD2 antibody ch14.18/CHO show higher ADCC levels and improved event-free survival. Oncoimmunology 5(11):e1235108.

30. Ott PA, et al. (2017) An immunogenic personal neoantigen vaccine for patients with melanoma. Nature 547(7662):217–221.

31. van Bergen J, et al. (2013) HLA reduces killer cell Ig-like receptor expression level and frequency in a humanized mouse model. J Immunol 190(6):2880–2885.

32. Cribbs A, et al. (2018) Inhibition of histone H3K27 demethylases selectively modulates inflammatory phenotypes of natural killer cells. J Biol Chem 293(7):2422–2437.

33. LaMere SA, Thompson RC, Komori HK, Mark A, & Salomon DR (2016) Promoter H3K4 methylation dynamically reinforces activation-induced pathways in human CD4 T cells. Genes Immun 17(5):283–297.

34. Kumar N & Khakoo SI (2018) Hepatocellular carcinoma: Prospects for natural killer cell immunotherapy. HLA.

35. Nakabayashi H, Taketa K, Miyano K, Yamane T, & Sato J (1982) Growth of human hepatoma cells lines with differentiated functions in chemically defined medium. Cancer Res 42(9):3858–3863.

36. Samson A, et al. (2018) Oncolytic reovirus as a combined antiviral and anti-tumour agent for the treatment of liver cancer. Gut 67(3):562–573.

37. Kurokohchi K, et al. (1996) Expression of HLA class I molecules and the transporter associated with antigen processing in hepatocellular carcinoma. Hepatology 23(5):1181–1188.

38. Zheng Y, et al. (2014) KPT-330 inhibitor of XPO1-mediated nuclear export has anti-proliferative activity in hepatocellular carcinoma. Cancer Chemother Pharmacol 74(3):487–495.

39. Kim J, et al. (2016) XPO1-dependent nuclear export is a druggable vulnerability in KRAS-mutant lung cancer. Nature 538(7623):114–117.

40. Chari A, et al. (2019) Oral Selinexor-Dexamethasone for Triple-Class Refractory Multiple Myeloma. N Engl J Med 381(8):727–738.

41. Birnbaum DJ, Finetti P, Birnbaum D, Mamessier E, & Bertucci F (2019) XPO1 Expression Is a Poor-Prognosis Marker in Pancreatic Adenocarcinoma. J Clin Med 8(5).

42. Camus V, Miloudi H, Taly A, Sola B, & Jardin F (2017) XPO1 in B cell hematological malignancies: from recurrent somatic mutations to targeted therapy. J Hematol Oncol 10(1):47.

43. Chen L, et al. (2019) Prognostic roles of the transcriptional expression of exportins in hepatocellular carcinoma. Biosci Rep 39(8).

44. Di Marco M, et al. (2017) Unveiling the Peptide Motifs of HLA-C and HLA-G from Naturally Presented Peptides and Generation of Binding Prediction Matrices. J Immunol 199(8):2639–2651.

45. Hilton HG, et al. (2017) The Intergenic Recombinant HLA-B *46:01 Has a Distinctive Peptidome that Includes KIR2DL3 Ligands. Cell Rep 19(7):1394–1405.

46. Fadda L, et al. (2010) Peptide antagonism as a mechanism for NK cell activation. Proc Natl Acad Sci U S A 107(22):10160–10165.

47. Sun JC, Beilke JN, & Lanier LL (2009) Adaptive immune features of natural killer cells. Nature 457(7229):557–561.

48. Fogel LA, Sun MM, Geurs TL, Carayannopoulos LN, & French AR (2013) Markers of nonselective and specific NK cell activation. J Immunol 190(12):6269–6276.

49. Wang AY & Liu H (2019) The past, present, and future of CRM1/XPO1 inhibitors. Stem Cell Investig 6:6.

50. Glasner A, et al. (2012) Recognition and prevention of tumor metastasis by the NK receptor NKp46/NCR1. J Immunol 188(6):2509–2515.

51. David G, et al. (2009) Discrimination between the main activating and inhibitory killer cell immunoglobulin-like receptor positive natural killer cell subsets using newly characterized monoclonal antibodies. Immunology 128(2):172–184.

52. Robinson MD, McCarthy DJ, & Smyth GK (2010) edgeR: a Bioconductor package for differential expression analysis of digital gene expression data. Bioinformatics 26(1):139–140.

53. Wu D & Smyth GK (2012) Camera: a competitive gene set test accounting for inter-gene correlation. Nucleic Acids Res 40(17):e133.

54. Alhamdoosh M, et al. (2017) Easy and efficient ensemble gene set testing with EGSEA. F1000Res 6:2010.

55. Chen J, Bardes EE, Aronow BJ, & Jegga AG (2009) ToppGene Suite for gene list enrichment analysis and candidate gene prioritization. Nucleic Acids Res 37(Web Server issue):W305–311.

56. Croft NP, et al. (2013) Kinetics of antigen expression and epitope presentation during virus infection. PLoS Pathog 9(1):e1003129.

57. Illing PT, et al. (2012) Immune self-reactivity triggered by drug-modified HLA-peptide repertoire. Nature 486(7404):554–558.

58. Schittenhelm RB, Sivaneswaran S, Lim Kam Sian TC, Croft NP, & Purcell AW (2016) Human Leukocyte Antigen (HLA) B27 Allotype-Specific Binding and Candidate Arthritogenic Peptides Revealed through Heuristic Clustering of Data-independent Acquisition Mass Spectrometry (DIA-MS) Data. Mol Cell Proteomics 15(6):1867–1876.

